# MicroRNA 1253 regulation of WAVE2 and its relevance to health disparities in hypertension

**DOI:** 10.1101/833673

**Authors:** Mercy A. Arkorful, Nicole Noren Hooten, Yongqing Zhang, Amirah N. Hewitt, Lori Barrientos Sanchez, Michele K. Evans, Douglas F. Dluzen

## Abstract

The prevalence of hypertension among African Americans (AAs) in the US is among the highest of any demographic and affects over two-thirds of AA women. Previous data from our laboratory suggests substantial differential gene expression (DGE) of mRNAs and microRNAs (miRNAs) exists within peripheral blood mononuclear cells (PBMCs) isolated from AA and white women with or without hypertension. We hypothesized that DGE by race may contribute to racial differences in hypertension. We found that the Wiskott-Aldrich syndrome protein Verprolin homologous-2 (*WAVE2*) is differentially-expressed in AA women with hypertension, along with several other members of the actin cytoskeleton signaling pathway that plays a role in cell shape and branching of actin filaments. We performed an *in silico* miRNA target prediction analysis that suggested miRNA miR-1253 regulates WAVE2. Transfection of miR-1253 mimics into human umbilical vein endothelial cells (HUVECs) and human aortic endothelial cells (HAECs) significantly repressed WAVE2 mRNA and protein levels (*P*<0.05), and a luciferase reporter assay confirmed that miR-1253 regulates the *WAVE2* 3’ UTR (*P*<0.01). miR-1253 over-expression in HUVECs significantly increased HUVEC lamellipodia formation (*P*<0.01), suggesting the miR-1253/WAVE2 interaction may play a role in endothelial cell shape and actin cytoskeleton function. Together, we have identified novel roles for miR-1253 and WAVE2 in a hypertension-related disparities context. This may ultimately lead to the discovery of additional actin-related genes which are important in the vascular-related complications of hypertension and influence the disproportionate susceptibility to hypertension among AAs in general and AA women in particular.

## Introduction

Throughout the United States, systemic arterial hypertension and hypertension-related conditions, including coronary atherosclerotic heart disease and cerebrovascular disease, have disproportionate incidence, mortality, and morbidity among African Americans (AAs). AA women are at particular risk. Between 2013-2016, 66% of AA females over ≥20 yrs had hypertension, compared with 41.3% of non-Hispanic white women, 41% of Hispanic women, and 36% of Asian women [1]. Reducing or eliminating hypertension is predicted to reduce cardiovascular disease (CVD)-related mortality in women by almost 40% [1, 2]. While 75% of AA women are aware of having hypertension, only 26% of AA women were able to control their high blood pressure [1]. A deeper understanding of the underlying biological mechanisms associated with hypertension may help reduce the burden of this condition.

Differential gene expression (DGE) can be linked with ancestry and can influence how individuals respond to environmental stimuli and exposures [3, 4] and their susceptibility to chronic diseases, including cancer [5] and peripheral arterial disease (PAD) [6]. Investigations have shown that DGE can predict outcomes to medical procedures including heart transplants [7]. DGE patterns are also linked with sex and gender. We previously reported that there is substantial differential mRNA and microRNA expression of hypertension-related genes and pathways in peripheral blood mononuclear cells (PBMCs) between AA and white women with hypertension [8, 9]. We observed that genes in canonical pathways related to hypertension, such as the renin-angiotensin (RAS) pathway, are expressed in reciprocal directions that is dependent upon race [9]. A follow-up analysis of these results identified that poly-(ADP-ribose) polymerase 1 (PARP-1), a DNA damage sensor protein involved in DNA repair and other cellular processes, is upregulated in hypertensive AA women compared with white hypertensive women and contributes to cellular response to inflammation [8]. AA women with PAD also have elevated levels of endothelial oxidative stress and circulating inflammatory biomarkers compared with AA men with PAD [6] and these differences may influence disease outcomes in AA women.

Understanding not only the significance of DGE patterns in hypertension and CVDs, but also the underlying genetic mechanisms that regulate these patterns, will help further our understanding of the biological basis of these conditions. Expression of hypertension-related genes can be regulated by ancestral genomic polymorphisms and expression quantitative train loci (eQTLs) [10, 11], but this does not account for all differences previously observed. This suggests alternative mechanisms also contribute to gene expression differences in different individuals. A possible contribution to variations in gene expression levels may arise from regulation from microRNAs.

MicroRNAs (miRNAs) are short (20-22 nucleotide), single-stranded RNAs that post-transcriptionally regulate protein expression by binding with target mRNA 3’ untranslated regions (UTRs) and inhibiting translation, often by degrading the target mRNA [12]. miRNA regulation of protein expression is integral to the proper functioning and health of the endothelial tissues of the vasculature, underlying smooth muscle layers, and vascular-response to changes in shear stress [13–15]. Disruption of miRNA regulation of hypertension-related genes can lead to endothelial dysfunction [14, 16, 17]. We previously reported that nine miRNAs exhibit disease-or race-specific differential expression and we have identified and validated novel hypertension-related targets for eight of these miRNAs [8, 9].

Here, we sought to identify and validate novel hypertension-related targets for miR-1253, which is significantly downregulated in PBMCs of hypertensive AA women [9], but had remained unexplored in our prior analyses. We have reanalyzed our microarray dataset to further our understanding of DGE in hypertensive women in hypertension-related pathways [9]. We identified significant DGE among genes within the actin-cytoskeleton signaling pathway between hypertensive AA and white women and we have validated hypertension-related miR-1253 as a novel regulator of WASP family Verprolin-homologous protein 2 (WAVE2), an integral member of the actin-cytoskeleton pathway.

## Materials and Methods

### Study Participants

Age-matched African American and white females who were either hypertensive (HT) or normotensive (NT) were previously chosen from the Healthy Aging in Neighborhoods of Diversity Across the Life Span (HANDLS) study of the National Institute on Aging Intramural Research Program (NIA IRP) of the National Institutes of Health (NIH) [18]. The demographics and clinical information for this re-examined cohort are previously described in extensive detail in [9]. The IRB of the National Institute of Environmental Health Studies, NIH, approved this study and all participants signed written informed consent.

### Microarray, Target Prediction, and Pathway Analysis

Gene expression levels in PMBCs reanalyzed in this study were analyzed and quantified using the Illumina Beadchip HT-12 v4 (San Diego, CA) as described in [9] and can be found in the Gene Expression Omnibus (<u>GSE75672</u>). Gene expression in HAECs was analyzed using the Illumina Beadchip HT-12 v4 and RNA was prepared and labeled according to the manufacturer’s protocol. Data were analyzed as previously performed [8] and outlying technical replicates were removed. Raw signals were analyzed by Z-score normalization [19] and individual genes with an average intensity >0, false discovery rate <0.2, *P*-value <0.05, and fold change >|1.5| were considered significant and these HAEC microarray datasets can be found in the Gene Expression Omnibus and will include our miR-1253 datasets (<u>GSE139286</u>). Gene expression data, including Z-ratio and fold-change, were imported into Ingenuity Pathway Analysis (IPA; Ingenuity Systems, Redwood City, CA) and we used default and custom settings to perform pathway analyses of genes significantly affected by miR-1253 over-expression and compared with a scrambled negative control. DIANA-Tarbase v7.0 [20] and TargetScan v7.2 [21] were used for miR-1253 target prediction.

### Cell Culture and Transfection

Primary human umbilical vein endothelial cells (HUVECs) were purchased and verified from Lonza and grown in EMB media supplemented with EGM-SingleQuot Kits (Lonza; Walkersville, MD). Primary human aortic endothelial cells (HAECs) were purchased from Lonza and grown in EMB-2 media supplemented with EGM-2 SingleQuot Kits (Lonza). Cells were transfected with miR-1253 Pre-miR miRNA Precursor (Assay ID #PM13220) or scrambled Pre-miR miRNA Precursor negative control #1 (Catalog #AM17110) (ThermoFisher, Waltham, MA). Mimics were transfected with Lipofectamine 2000 (ThermoFisher).

### 3’ UTR Luciferase reporter assays

Two miTarget miRNA 3’ UTR plasmids were purchased from GenoCopeia (Rockville, MD) containing either the first (Catolog #Hmi088372a-MT06) or second halves (Catalog #Hmi088372b-MT06) of the *WAVE2* 3’ UTR RNA sequencing. The miTarget plasmid vector (pEZX-MTO6) contains a luciferase reporter gene with attached 3’ UTRs of interest and downstream renilla luciferase for transfection efficiency controls. HUVECs were co-transfected with 50 ng of either WAVE2 3’UTR plasmid and with either 50 nM scrambled negative control or miR-1253 mimic. Forty-eight hours later, luciferase and renilla activities were measured using the Dual-Luciferase reporter assay system (Promega) according to manufacturer’s instructions. Renilla served as an internal transfection control and the ratio of luciferase/renilla was normalized to the scrambled control. All luciferase assays were measured using a Synergy HT Microplate Reader (BioTek, Winooski, VT) and performed in triplicate.

### RNA Isolation and RT-qPCR

Total RNA was isolated from HAECs and HUVECs using Trizol Reagents (ThermoFisher) with phenol/chloroform extraction according to manufacturer’s protocol. RNA integrity was measured with a Nanodrop 2000 and cDNA was synthesized using random hexamers and Super Script II reverse transcriptase (Invitrogen, Carlsbad, CA). miRNA cDNA was synthesized using the QuantiMiR RT Kit and the provided universal reverse primer (Systems Biosciences, Mountain View, CA). All RT-qPCR reactions were performed with 2x SYBR green master mix (ThermoFisher) on either an Applied Biosystems model 7500 real-time PCR machine or a QuantStudio 6 Flex. miR-1253 levels were normalized to *U6* and *WAVE2* levels were normalized to the average of *GAPDH* and *ACTB*. The following primers (forward and reverse) were used for each gene: miR-1253 forward 5’-AGAGAAGAAGATCAGCCTGCA-3’; *U6* forward 5’-CGCAAGGATGACACGCAAATTC-3’; *WAVE2* forward 5’– GCAGCATTGGCTGTGTTGAA-3’and reverse 5’-GGTTGTCCACTGGGTAACTGA-3’; *ACTB* forward 5’-GGACTTCGAGCAAGAGATGG-3’ and reverse 5’-AGCACTGTGTTGGCGTACAG-3’; *GAPDH* forward 5’-GCTCCTCCTGTTCGACAGTCA-3’ and reverse 5’-ACCTTCCCCATGGTGTCTGA-3’. Gene expression levels were calculated using the 2^−ΔΔCt^ methodology [22].

### Western blot analysis

HAECs and HUVECs were washed 2x with cold PBS and then lysed in 2x Laemmli sample buffer on ice. Protein lysate was then loaded into a 10% polyacrylamide gel and separated. Protein levels were determined by anti-WAVE2 (sc-373889; Santa Cruz Biotechnology, Dallas, TX), anti-GAPDH (c-32233; Santa Cruz), and anti-ACTB (sc-1616; Santa Cruz) antibodies. Densitometry was performed using ImageJ software [23].

### Immunofluorescence and Scoring of Cells with Lamellipodia and Filopodia

HUVECs were fixed in formaldehyde on glass slides and permeabilized in Triton-X. Cells were stained with Rhodamine Phalloidin (1:300) (Life Technologies), then with DAPI (1:10,000) and then mounted using ProLong (ThermoFisher Scientific). HUVECs were scored positive for the presence of lamellipodia if they displayed at least one actin-rich (phalloidin positive) ruffled structure at the edge of the cell. Filopodia were scored positive if at least two actin-positive finger-like protrusions were observed emanating from the cell. We used a Zeiss Observer D1 microscope with an AxioCam1Cc1 camera. Only cells that were either isolated or only attached to one other cell were counted. The number of positive cells is shown as a ratio to all DAPI-stained cells and cell area was measured using AxioVision Rel 4.7 software. This approach was modified from [24].

### Statistical Analysis

The Student’s *t*-test was used when comparing two groups unless otherwise indicated. A *p*-value of <0.05 was considered statistically significant and calculations were performed in Prism GraphPad v8.2.0, unless otherwise indicated.

## Result

We used the DIANA-Tarbase v7.0 [20] and TargetScan v7.2 [21] algorithms to identify potential miR-1253 mRNA targets in humans. DIANA-Tarbase predicted 4,723 mRNAs as potential targets and TargetScan identified 5,345 mRNAs (Figure 1A, see Supp. File 1 for complete list). There were 2,885 unique mRNAs that overlapped between both prediction programs and we used this list moving forward with our *in silico* analysis. We compared the 2,885 mRNAs with the 3,354 mRNAs found to be differentially-expressed in PBMCs in our hypertension cohort when comparing gene expression between AA and white women with or without hypertension [9]. We found that 840 of the miR-1253 predicted targets exhibited differential-expression in PBMCs (Figure 1B; Supp. File 1) and 112 of these predicted targets are also found in our previously-curated list of 1,266 genes related to hypertension and inflammation (Figure 1C; Supp. File 1) [9].

**Figure 1:**
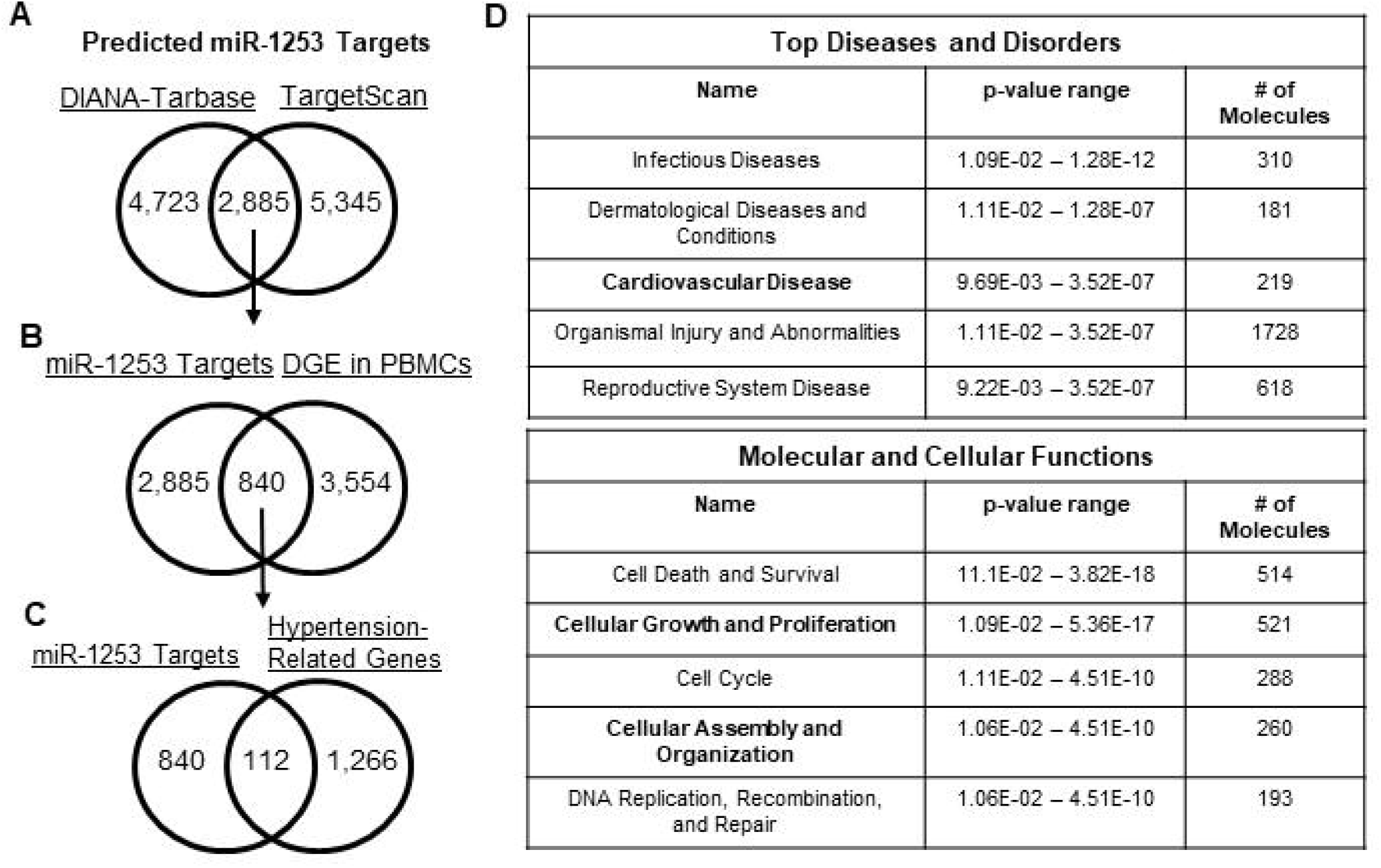
Target prediction analysis for miR-1253. **(A)** Venn diagram of miR-1253 predicted mRNA targets overlapping between the DIANA-Tarbase and TargetScan algorithms. **(B)** Venn diagram of overlapping, predicted miR-1253 targets that are differentially expressed in PBMCs identified in [9]. **(C)** Venn diagram of miR-1253 predicted targets that are significantly, differentially-expressed in PBMCs and within hypertension-related pathways identified using Ingenuity Pathway Analysis (IPA). **(D)** List of significant Top Diseases and Disorders (Top) and Molecular and Cellular Functions (Bottom) in HAECs transfected with 50 nM miR-1253 mimic. These pathways were identified by Ingenuity Pathway Analysis.

We next sought to further parse down this list of 112 mRNA targets and validate the role of miR-1253 in potentially regulating expression of some of these mRNAs. We over-expressed 50 nM of miR-1253 mimic in human aortic endothelial cells (HAECs) for 48 hours and performed a discovery microarray to assess gene expression level changes. We used Ingenuity Pathway Analysis (IPA) to identify the Top Disease and Disorders and Molecular and Cellular Functions associated with miR-1253 over-expression. We observed that pathways related to cardiovascular disease, cellular growth and proliferation, and cellular assembly and organization were the most significantly affected in response to miR-1253 expression and within the top five of pathways in each category (Figure 1D).

We next examined DGE in the actin cytoskeleton signaling pathway in our hypertension cohort by reanalyzing our previous microarray dataset <u>GSE75672</u>. We chose this pathway given the role of actin cytoskeletal remodeling and signaling in hypertension and endothelial function [25–27] and the importance of this pathway in CVD and cellular growth and proliferation identified in IPA (Figure 1D). We used IPA to overlay mRNA expression in PBMCs that were isolated from 24 age-matched females who were either African American normotensive women (AANT), African American hypertensive women (AAHT), white normotensive women (WNT), or white hypertensive women (WHT; n=6/group, as previously extensively described in [9]) to identify DGE in the actin cytoskeleton signaling pathway.

While only *PAK* was significantly higher in AANT compared with WNT in this pathway (Figure 2A), we found that 27 genes of the 75 in the actin cytoskeleton signaling pathway are significantly different (*P*<0.05 and |fold-change| >1.5; Supp. Table 1) when comparing AAHT with WHT (Figure 2B). There are only three genes significantly different in this pathway between WHT and WNT in our cohort (*ARP2*, *ACTG1 [F-actin]*, and *SRC;* Supp. Figure 1A) and *ARP2* and *ACTG1* are reciprocally-expressed when comparing AAHT with AANT (Supp. Figure 1B), suggesting that these genes exhibit DGE by race in hypertensive women. We also observed that there are more genes significantly different when comparing AAHT with AANT (Supp. Figure 1B) than when comparing WHT with WNT, suggesting that the actin cytoskeleton signaling pathway is an overlooked gene pathway when examining health disparities in hypertension, particularly in AA women.

**Figure 2:**
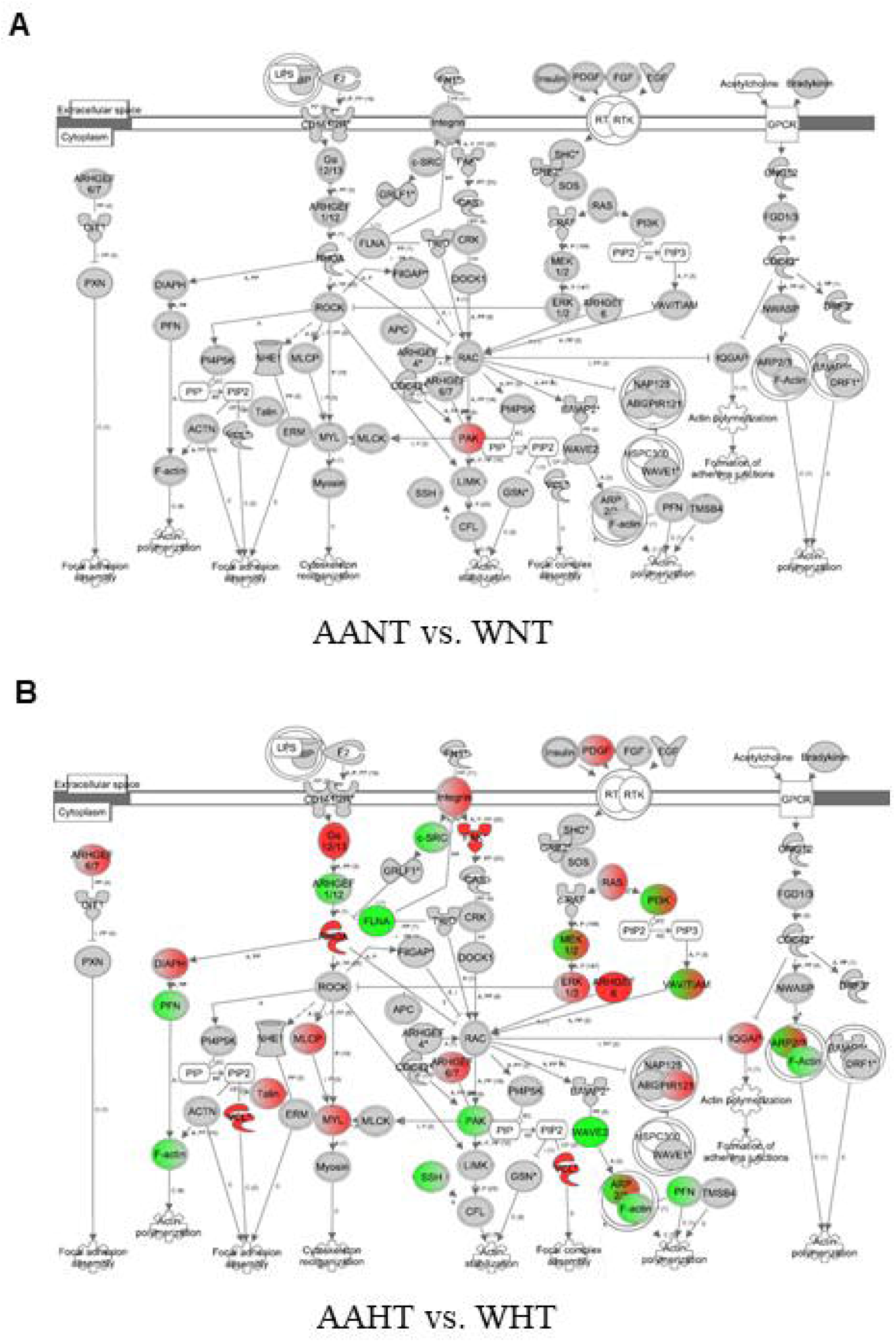
Gene expression analysis of the actin cytoskeleton in hypertensive women. Microarray gene expression fold-changes in PBMCs isolated from AANT, WNT, AAHT, and WHT were imported into Ingenuity Pathway Analysis (IPA) and overlaid onto the actin cytoskeleton pathway. Red indicates significantly up-regulated expression and green indicates significant down-regulation in AANT compared with WNT **(A)** and in AAHT compared with WHT **(B)**. Grey indicates a non-significant difference and white indicates no data available. All fold changes and P-values are listed for each gene and each comparison in Supplementary Table 1. AANT: African American normotensive women; AAHT: African American hypertensive women; WNT: White normotensive women; WHT: White hypertensive women.

In order to determine whether miR-1253 might play a role in the differential-expression of genes within the actin cytoskeleton signaling pathway, we compared those mRNAs significantly down-regulated in HAECs via over-expression of miR-1253 mimic against the 1,266 genes in our hypertension gene list. There were 747 mRNAs significantly repressed >1.5-fold compared with the scrambled negative control (*P*<0.05; FDR <0.20; n=5; Supp. File 1). Of these 747, 23 mRNAs are within our hypertension gene list and significantly different in our hypertension cohort (Table 1). One of these genes, WASP family Verprolin-homologous protein 2 (WAVE2), plays a role in the regulation of the actin cytoskeleton [24, 28]. miR-1253 is also predicted to target two other genes in the actin cytoskeleton pathway, Filamin A, Alpha (*FLNA)* and Ras Homolog A (*RHOA)*, however, neither of these two mRNAs were significantly down-regulated by miR-1253 in our screen. Therefore, we focused on *WAVE2* as a potential target of miR-1253.

**Table 1:** Summary of miR-1253 Predict Targets Repressed in HAECs*.

| Predicted miR-1253 Targets (compared w/scrambled control) | FDR | Fold Change | P-Value | Z-Ratio |
| --- | --- | --- | --- | --- |
| ABCB10 | 0.0172 | -1.53 | 0.0032 | -2.83 |
| ACO1 | 0 | -7.58 | 0 | -11.94 |
| ACSL1 | 0 | -1.69 | 0 | -3.32 |
| DCUN1D5 | 0.0116 | -1.87 | 0.002 | -3.87 |
| DPYSL2 | 0 | -1.61 | 0 | -2.47 |
| DUSP14 | 0 | -2.13 | 0 | -4.46 |
| MSN | 0 | -1.83 | 0 | -3.15 |
| PARP1 | 0 | -4.4 | 0 | -8.57 |
| PDE12 | 0 | -2 | 0 | -4.36 |
| POLA1 | 0 | -1.53 | 0 | -2.82 |
| PTGER4 | 0 | -1.72 | 0 | -3.05 |
| RAB27A | 0 | -2.06 | 0 | -4.5 |
| RSU1 | 0 | -2.16 | 0 | -4.78 |
| RXRA | 0 | -1.54 | 0 | -2.61 |
| SEC62 | 0 | -1.55 | 0 | -2.96 |
| SERINC3 | 0 | -2.51 | 0 | -5.53 |
| SPARC | 0 | -1.91 | 0 | -3.55 |
| TFRC | 0 | -1.71 | 0 | -3.1 |
| TMEM127 | 0 | -1.89 | 0 | -4.07 |
| TNS3 | 0 | -1.83 | 0 | -3.68 |
| TOPBP1 | 0 | -2.09 | 0 | -4.35 |
| UBE2N | 0.0005 | -1.56 | 0.0001 | -2.67 |
| WAVE2 | 0 | -1.64 | 0 | -3.09 |
| *mRNAs also found to be significantly and differentially expressed in PMBCs in hypertensive AA and white women and with hypertension related pathways outlined in [9]. |  |  |  |  |

We performed a luciferase reporter assay using miTarget reporter vectors to confirm that miR-1253 can regulate the 3’ untranslated region (UTR) of *WAVE2*. The 3’ UTR of *WAVE2* is 3,959 nucleotides in length and was split between two miRTarget plasmids. These heterologous reporter plasmids contain luciferase with a downstream renilla luciferase (RL) transfection control. The miRTarget *WAVE2* 3’UTR-1 plasmid contains the first 2,010 nucleotides of the *WAVE2* 3’ UTR, including the last 21 nucleotides of its coding region. The miRTarget *WAVE2* 3’ UTR-2 plasmid contains nucleotides 1,888 to 3,959 of the *WAVE2* 3’ UTR and there is a common overlap of 122 nucleotides of the 3’UTR between plasmid 1 and 2 (Figure 3A). TargetScan predicted that miR-1253 binds to the *WAVE2* 3’ UTR at nucleotides 3,734 to 3,756 in the second half of the *WAVE2* 3’ UTR, which is referred to as the *WAVE2* 3’UTR binding site #3 (Figure 3B). We also identified potential seed region binding sites at two additional positions at nucleotides 1,617 to 1,622 (binding site #1) and 1,775 to 1,780 (binding site #2), which are found in the first half of the 3’UTR (Figure 3A). Human umbilical vein endothelial cells (HUVECs) were co-transfected with 50 nM miR-1253 or scrambled control mimics and either miRTarget *WAVE2* 3’UTR-1 or 3’UTR-2. We observed significant repression of luciferase activity for miRTarget 3’UTR-1 (*P*<0.01, n=3) and miRTarget 3’UTR-2 (*P*<0.001, n=3) in the presence of miR-1253 and compared to scrambled control (Figure 3C). These data indicated that miR-1253 can bind to the *WAVE2* 3’ UTR and reduce protein expression.

**Figure 3:**
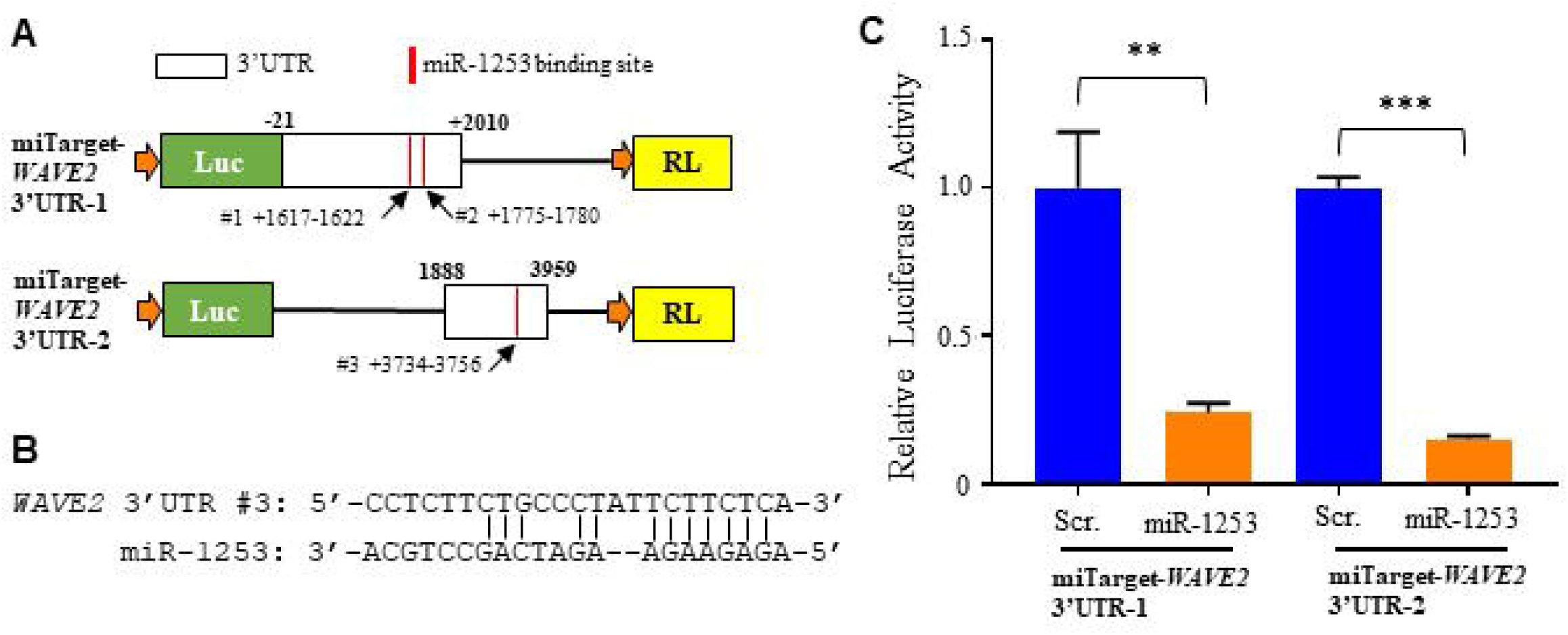
miR-1253 targeting of the *WAVE2* 3’UTR. **(A)** Schematic of the miRTarget *WAVE2* 3’ UTR vectors (plasmid 1 and 2). The predicted binding sites of miR-1253 to *WAVE2* 3’UTR are indicated in red with designated base pair positions. **(B)** Base pair schematic of binding site #3 of miR-1253 to the 3’ UTR region of WAVE2 as predicted by TargetScan. **(C)** The relative expression of luciferase (Luc) reporter in the presence of 50 nM miR-1253 for 48 hrs and compared with scrambled control. Data were normalized to an internal renilla control and normalized to 1.0. \*\**P*<0.01; **P<0.001, by two-tailed student’s T-test.

We next validated whether miR-1253 can regulate WAVE2 expression *in vitro*. We over-expressed 50 nM miR-1253 mimic for 48 hours in human aortic endothelial cells (HAECs). In the presence of miR-1253, *WAVE2* mRNA levels were significantly repressed nearly 50% (*P*<0.05; n=3) and the corresponding WAVE2 protein levels were significantly down-regulated by nearly 60% (*P*<0.01; n=3) compared with a scrambled control mimic (Figure 4A). In order to verify this is not a cell-line specific effect, we also performed the same experiments in HUVEC cells. miR-1253 mimics significantly repressed *WAVE2* mRNA 55% (*P*<0.01; n=5) and WAVE2 protein 38% (*P*<0.001; n=5) (Figure 4B). Together, these results confirm our *in silico* prediction that miR-1253 can regulate the expression of WAVE2 protein in endothelial cells.

**Figure 4:**
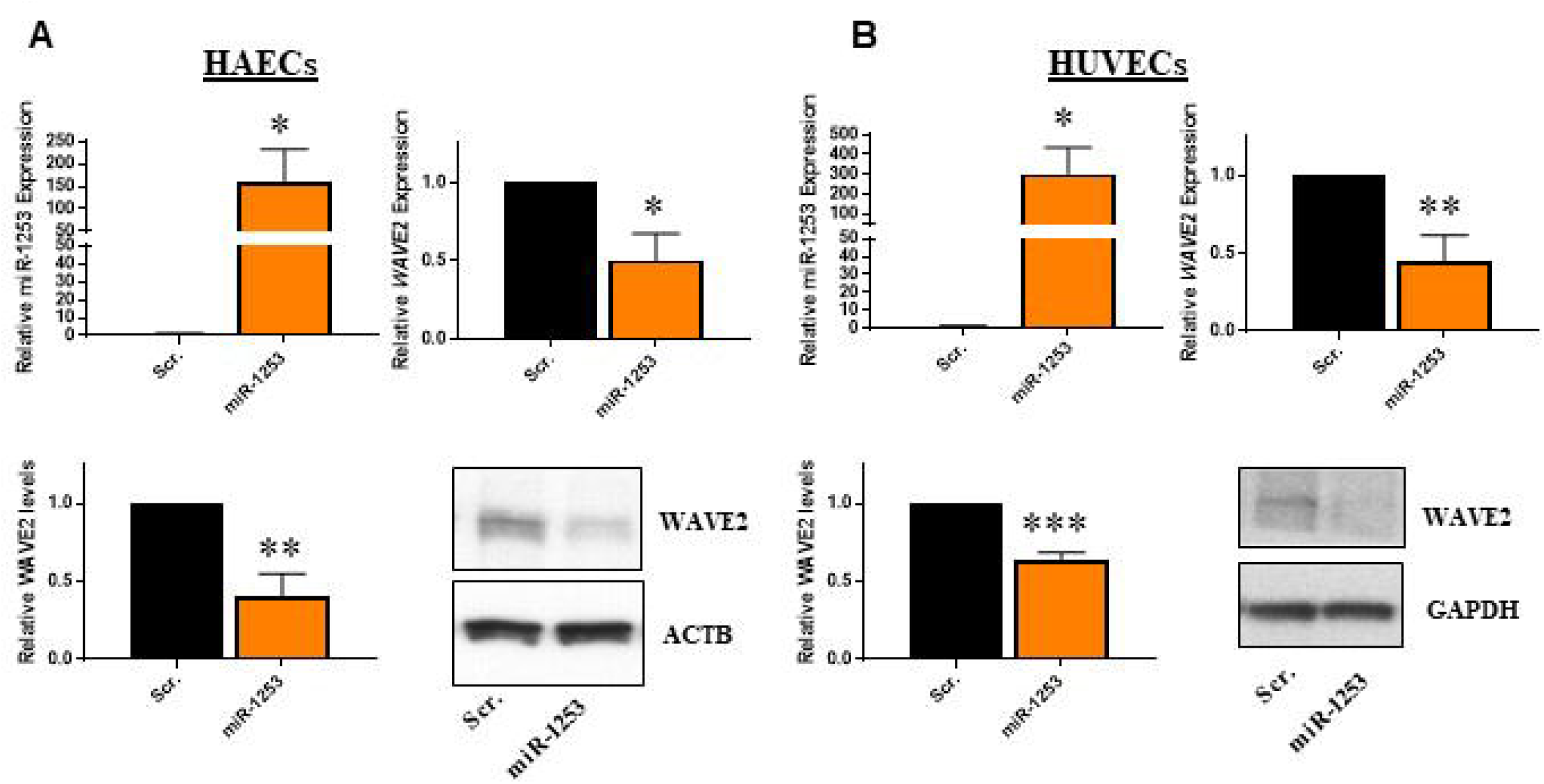
Overexpression of miR-1253 in HAECs and HUVECs reduces expression of WAVE2. 50 nM miR-1253 was transfected into HAECs (n=3) **(A)** and HUVECs (n=5) **(B)** for 48 hrs and over-expressed in each cell line compared with a scrambled negative control mimic (scr.; top left). *WAVE2* expression was normalized to *GAPDH* in each cell line and shown relative to scrambled (scr.; top right). WAVE2 proteins levels were normalized to Beta Actin (HAECs) or GAPDH (HUVECs) and shown relative to a scrambled control (scr.; bottom left). Representative immunoblots are shown for WAVE2 and loading controls in each cell line (bottom right) \**P*<0.05, \*\**P*<0.01, \*\*\**P*<0.001, by one-tailed T-test (for confirmation of miR-1253 expression levels in each cell line) or two-tailed student’s T-test for all comparisons of mRNA and protein levels.

Given that WAVE2 is a key regulator of actin cytoskeleton dynamics, we assessed whether this regulatory network may affect the actin cytoskeleton. We transfected 50 nM scrambled control or miR-1253 mimics into HUVECs for 48 hours and stained with rhodamine phalloidin to visualize actin cytoskeletal structures. We observed morphological changes in cells transfected with miR-1253 mimic compared to scrambled control mimics (Figure 5). Protrusive actin-containing structures such as lamellipodia or filopodia are formed at the leading edge of cells. Lamellipodia form larger actin-containing ruffles while filopodia are characterized by actin-containing finger-like extensions from the cell. Cells with transfected miR-1253 had increased lamellipodia formation as shown by concentrated actin-rich membrane-ruffling at the edges of cells. Therefore, we scored these cells by the presence of either lamellipodia or filopodia. We observed that there was a significant increase in lamellipodia in HUVECs with miR-1253, indicating an increase in actin-rich membrane ruffling at the edges of the cells (*P*<0.001; n=3) (Figure 5A and 5B). miR-1253 did not affect the formation of actin-rich filopodia projections. We did observe an increase in cell surface area of approximately 60%, however this was not statistically significant (*P*=0.09; n=3) (Figure 5C). Together, miR-1253 regulate *WAVE2* in endothelial cells leading to changes in endothelial cell lamellipodia formation.

**Figure 5:**
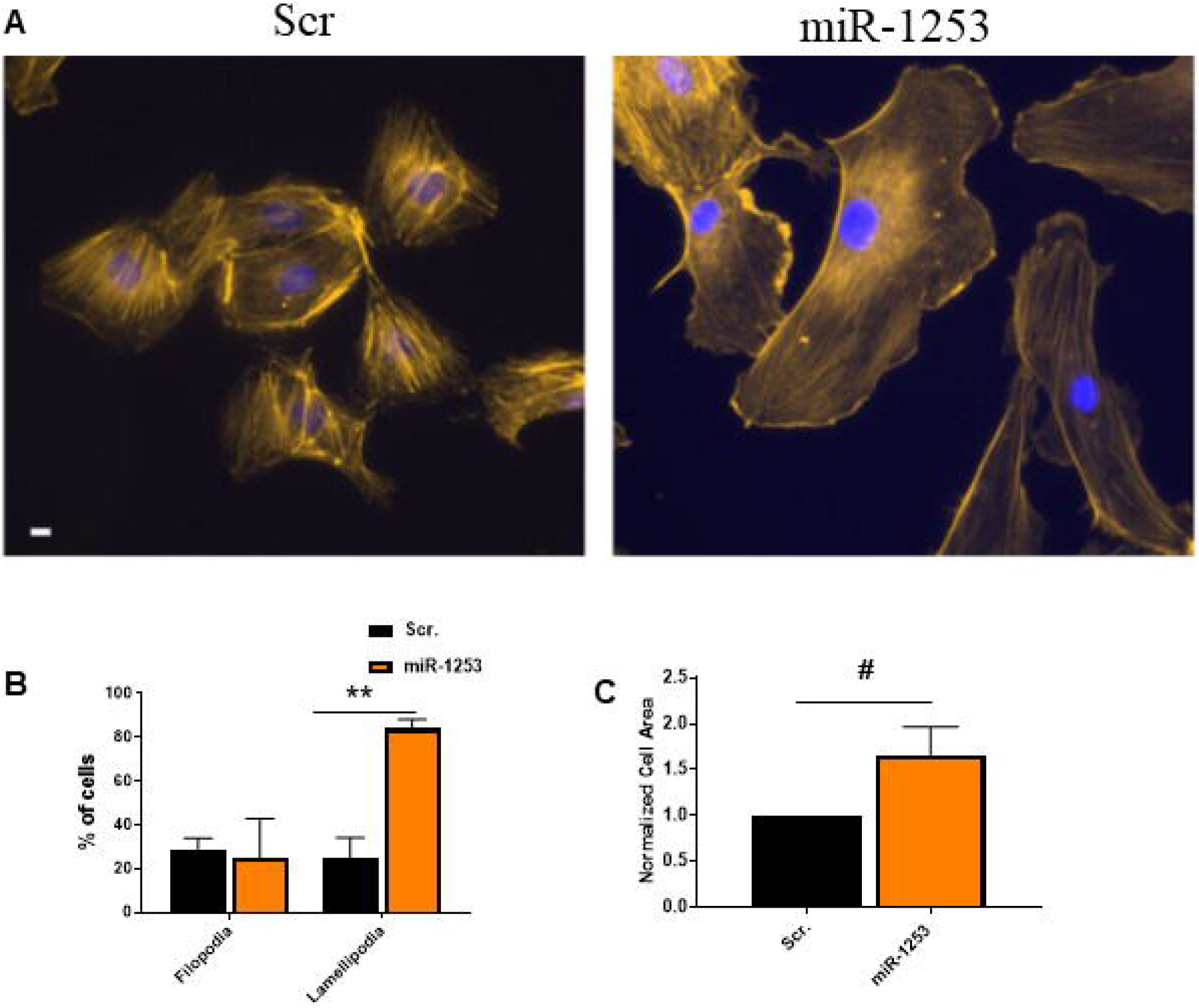
miR-1253 increases lamellipodia in HUVECs. **(A)** Representative pictures of HUVECs transfected with either scrambled control mimics (scr., left panel) or miR-1253 (right panel) and stained with rhodamine phalloidin for actin filaments and DAPI for nuclei visualization. **(B)** Percent of cells visualized and counted for filopodia or lamellipodia in cells transfected with the scrambled or miR-1253 mimic versus total number of DAPI-stained cells (n=3). (C) Quantitation of cell surface area of HUVECs transfected with the scr. control or miR-1253 mimic (n=3). \*\**P*<0.01, #*P*=0.09; Two-tailed student’s T-test. Scale bar = 10 µm.

## Discussion

Together, our data indicate that a large number of genes within the actin cytoskeleton signaling pathway are differentially-expressed in PBMCs between AA and white hypertensive women, with nearly all of these genes exhibiting similar expression levels between normotensive AA and white women (Figure 2, Supp. Fig 1). This suggests that the DGE patterns associated with hypertension occur sometime as the disease process begins or after sustained exposure to elevated systemic blood pressure levels. Previously, we found similar patterns in additional pathways related to hypertension [8, 9] providing further evidence that DGE is associated with individual gene expression levels in individuals with high blood pressure. Here, we found that miR-1253, identified in our previous analysis [9] but without a functionally-validated role in hypertension, was predicted to target *WAVE2* in the actin cytoskeleton pathway. This gene expression analysis in PBMCs led us to validate that miR-1253 can bind and regulate WAVE2 expression in endothelial cells and influence actin cytoskeletal dynamic (Figures 4-5).

DGE within the actin cytoskeleton signaling pathway in hypertension has previously remained relatively unexplored, particularly in the context of AA women with hypertension. Most studies have examined the role of this pathway in downstream conditions of which hypertension is a major risk factor. Pathway analysis of gene expression in coronary artery atherosclerosis plagues identified that focal adhesion and actin cytoskeleton pathways as some of the most differentially-expressed between early and late-stage plaques [29]. In human macrophages, *FLNA* expression is higher in advanced atherosclerotic plaques compared with intermediary plaques and inhibition of *FLNA* expression in mice reduced plaque development, suggesting a role for this gene and the actin cytoskeleton in hypertension-related CVDs [30]. *FLNA* is expressed in human and mouse endothelial cells after myocardial infarction. When its expression is inhibited the endothelial response to cardiac repair, migration, and VEGF-A secretion was reduced and this promoted left ventricular dysfunction and heart failure [31]. In our analysis, we found that AA women have higher levels of *FLNA* compared with white women which may suggest that additional members of this pathway are relevant in hypertension etiology and the development of end organ complications.

Altered levels of other members of the actin cytoskeleton signaling pathway have been observed but not in the context of gender or race. Bradykinin receptors 1 and 2, which act as upstream regulators of vessel wall remodeling, are significantly upregulated in peripheral monocytes of essential hypertensives and hypertension treatment reduces their expression [32]. The Rho/ROCK signaling cascade regulates organization of the actin cytoskeleton and cell morphology, including adhesion of cells along the endothelium of the vasculature [33–35]. Members of the RhoA family have been extensively examined as targets for hypertension therapy [36], and given its upregulation in AA women with hypertension [shown here and in [9]], the targeting of elevated *RHOA* expression and the downstream impact on cytoskeleton function may be a novel area for intervention in AA women. Follow-up studies are warranted to investigate this.

We identified differential expression of *WAVE2* between AA and white women with hypertension. WAVE2 is an actin nucleation promoting factor and binds with the actin-related protein (Arp) 2/3 complex to promote actin filament nucleation and branching [37, 38]. Variation in WAVE2 expression modulates actin branching and influences the formation of cellular filopodia and lamellipodia [24, 28, 38–40]. We observed that repression of *WAVE2* levels due to miR-1253 overexpression increased the formation of lamellipodia and membrane ruffling, consistent with lamellipodia formation and actin elongation dynamics related to WAVE2 expression modulation [28]. It’s possible that miR-1253 regulation of *WAVE2* in hypertensives may influence endothelial integrity and lead to downstream complications and additional studies are warranted to investigate this.

Modification of WAVE2 expression by miR-1253 in either circulating monocytes or in endothelial cells may be associated with hypertension-related changes in membrane physiology and morphology. Endothelial response to increase shear stress and laminar flow has been found to be associated with race. HUVECs isolated from AAs are more responsive to laminar shear stress compared with HUVECs from whites, including in pathways related to nitric oxide synthase and oxidative stress response. Importantly, in both cases, exercise was able to improve upon those changes [41, 42]. A recent meta-analysis of studies comparing arterial stiffness between AAs and whites identified significant differences in AAs in aortic femoral pulse wave velocity and carotid-femoral pulse wave velocity [43] and build off of previous analysis that AAs can have impaired microvascular dilatory response [44]. Our findings here indicate that the actin cytoskeleton could influence or associate with these clinical observations and further consideration of the involvement of WAVE2, miR-1253, and related pathway genes will be important to identify any direct roles.

Our analysis identifies a novel regulatory role for miR-1253 and regulation of WAVE2. Previously, the only known role for miR-1253 has been found in cancer, where it regulates the expression of the long, non-coding RNA *FOXC2-AS1* in prostate cancer cells [45] and *WNT5A* in lung carcinoma [46]. This study is limited because it is not known whether differential-expression of miR-1253 in AA women with hypertension is a contributing cause or an effect of elevated high blood pressure. There is no data in the literature examining whether changes in miR-1253 influence endothelial dysfunction via changes to the actin cytoskeleton and if these changes predispose individuals to atherosclerotic plaques or other hypertension-related complications. These questions were beyond the scope of this study. This finding also underscores the need to validate if miR-1253 regulates WAVE2 expression in vascular smooth muscle cells, as this tissue is linked with the endothelial layer of the vasculature, or if miR-1253 expression changes in response to shear stress.

Many miRNAs play an important role in the normal and disease physiology of the vasculature. For example, miR-155 regulates endothelial *eNOS* and downstream vasodilation in human mammary arteries [47] and its expression is inversely correlated with target *AGTR1* expression in untreated hypertensives [48]. Several miRNAs, including miR-143 and miR-145, regulate vascular smooth muscle cell function and have been found to be differentially-expressed in PBMCs and correlated with 24-hr diastolic blood pressure and pulse pressure in individuals with hypertension [49]. It unknown if these miRNAs are correlated with disparities in hypertension, particularly in AA women, or involved in similar pathways as miR-1253.

Together, we have identified the actin cytoskeleton as a possible avenue to explore to further our understanding of how hypertension may develop and present in different populations. Importantly, we have identified a bioinformatic analysis pipeline that can identify and validate novel miRNA regulators for members of that pathway. Future studies will need to examine the *WAVE2*-miR-1253 relationship in order to further elucidate their role in hypertension and hypertension-related disparities.

## Supporting information

Supplemental Gene Lists

## Supplementary Materials

Supplementary Table 1 and Supplementary Figure 1 can be found in the Supplementary Materials document. All gene lists from our miR-1253 prediction analysis, microarray analysis, and endothelial cells studies can be found in the excel file labeled Supplemental File 1.

## Acknowledgements

We thank Dr. Simonetta Camandola for the use of the luminometer for the luciferase assay readings and Dr. Elin Lehrmann for assistance with our microarray. We thank Althaf Lohani for preparing materials for the luciferase assays and for RNA isolations. We thank the HANDLS staff for their critical evaluation of HANDLS participants.

## Author Contributions

Conceptualization, M.A.A., N.N.H., M.K.E., and D.F.D. Methodology and Validation, M.A.A., N.N.H., Y.Z., A.N.H., L.B.S., D.F.D. Formal Analysis,: M.A.A, N.N.H., Y.Z., A.N.H, L.B.B., M.K.E., and D.F.D. Writing Draft Preparation and Editing: M.A.A., N.N.H., and D.F.D. Contributed reagents/resources/software: N.N.H., Y.Z., M.K.E., and D.F.D. All authors reviewed the final manuscript.

## Data Availability Statement

The datasets analyzed for this study can be found in the Gene Expression Omnibus (<u>GSE75672</u>) and (<u>GSE139286</u>).

