## Supplemental Table 1 and Figure 1 for "MicroRNA 1253 regulation of WAVE2 and its relevance to health disparities in hypertension"

Supplementary Material

**Supplementary Table 1 – Genes within the Actin Cytoskeleton Signaling Pathway with Corresponding Fold Change and P-Value**

| **Actin Cytoskeleton** | **(Fold Change) AAHT-AANT** | **(p) AAHT-AANT** | **(Fold Change) AAHT-WHT** | **(p) AAHT-WHT** | **(Fold Change) AANT-WNT** | **(p) AANT-WNT** | **(Fold Change) WHT-WNT** | **(p) WHT-WNT** |
| --- | --- | --- | --- | --- | --- | --- | --- | --- |
| **ABI2** | -1.17 | 0.005 | -1.24 | 0.0174 | 1.26 | 3.82E-05 | -1.1 | 0.3265 |
| **ACTN** | -1.097 | 3.86E-01 | -1.095 | 3.68E-01 | -1.018 | 4.43E-01 | 1.016 | 4.97E-01 |
| **APC** | -1.011 | 8.58E-01 | -1.094 | 7.00E-01 | 1.135 | 7.72E-01 | -1.048 | 9.82E-01 |
| **ARHGEF1** | **-2.418** | **8.48E-04** | **-2.911** | **2.93E-04** | -1.164 | 8.02E-01 | 1.379 | 2.46E-01 |
| **ARHGEF12** | 1.188 | 1.47E-02 | 1.244 | 1.40E-04 | 1.093 | 8.08E-01 | -1.145 | 3.47E-02 |
| **ARHGEF4** | 1.09 | 0.0006 | -1.08 | 0.9989 | 1.19 | 0.5152 | -1.07 | 0.9037 |
| **ARHGEF6** | 1.36 | 0.0374 | **1.57** | **0.025** | -1.1 | 0.4827 | -1.05 | 0.248 |
| **ARHGEF7** | 1.121 | 8.93E-03 | 1.434 | 7.79E-03 | -1.198 | 3.71E-01 | -1.052 | 9.90E-01 |
| **ARP2 (ARPC4)** | **-1.566** | **4.99E-02** | **-2.189** | **1.11E-02** | -1.264 | 2.33E-01 | **1.767** | **3.75E-02** |
| **ARP3 (ACTR3)** | **1.553** | **3.75E-02** | **2.175** | **5.44E-04** | -1.28 | 4.66E-02 | -1.094 | 3.88E-01 |
| **BAIAP2** | -1.3 | 0.0025 | -1.24 | 0.082 | 1.18 | 0.2823 | -1.19 | 0.0544 |
| **Bradykinin** | 1.07 | 0.0986 | -1.02 | 0.1622 | 1.25 | 0.0334 | -1.15 | 0.0448 |
| **CAS** | 1.08 | 0.0411 | 1.02 | 0.0864 | 1.22 | 0.041 | -1.15 | 0.1168 |
| **CD14** | -1.53 | 0.2323 | 1.65 | 0.0684 | -1.45 | 0.6614 | 1.04 | 0.92 |
| **CDC42** | -1.28 | 0.0175 | -1.25 | 0.0736 | 1.18 | 0.4037 | -1.15 | 0.5953 |
| **CFL** | 1.057 | 8.82E-01 | 1.146 | 9.13E-01 | 1.036 | 1.87E-01 | -1.123 | 2.69E-01 |
| **c-RAF** | 1.34 | 0.079 | 1.34 | 0.0961 | -1.03 | 0.5384 | 1.03 | 0.5993 |
| **CRK** | -1.437 | 7.07E-03 | -1.392 | 1.06E-02 | -1.26 | 1.33E-01 | 1.217 | 7.59E-02 |
| **c-SRC** | **-1.715** | **5.60E-03** | **-1.529** | **5.87E-02** | 1.202 | 1.68E-01 | -1.158 | 1.26E-01 |
| **DIAPH** | **1.604** | **2.78E-03** | **2.038** | **6.43E-07** | 1.252 | 1.05E-01 | -1.171 | 8.46E-02 |
| **DOCK1** | 1.05 | 0.233 | 1.01 | 0.0378 | 1.19 | 0.4453 | -1.14 | 0.144 |
| **DRF1** | -1.33 | 0.0022 | -1.27 | 0.2419 | -1.03 | 0.8904 | 1.03 | 0.3888 |
| **DRF3** | -1.15 | 0.0075 | -1.18 | 0.0975 | 1.24 | 0.0503 | -1.14 | 0.0914 |
| **EGF** | -1.13 | 0.1564 | 1.07 | 0.0524 | 1.04 | 0.5801 | -1.26 | 0.0499 |
| **ERK1** | -1.105 | 4.08E-01 | -1.367 | 8.62E-02 | -1.025 | 8.60E-01 | 1.22 | 1.95E-01 |
| **ERK2** | 1.482 | 1.24E-02 | **1.748** | **5.69E-04** | -1.249 | 2.86E-02 | 1.182 | 2.17E-01 |
| **ERM (MSN)** | **-1.586** | **2.71E-07** | -1.337 | 5.62E-02 | -1.033 | 6.21E-01 | -1.149 | 2.92E-01 |
| **F2** | 1.1 | 0.1521 | 1.03 | 0.1626 | 1.14 | 0.6609 | -1.07 | 0.7739 |
| **F2R** | 1.19 | 0.1907 | 1.15 | 0.2103 | 1.15 | 0.1638 | -1.17 | 0.279 |
| **F-actin (ACTG1)** | **-1.731** | **7.97E-02** | **-2.748** | **5.33E-03** | -1.326 | 5.64E-01 | **2.105** | **3.79E-02** |
| **FAK** | 1.45 | 0.0902 | **2.13** | **0.0001** | 1.12 | 0.9882 | -1.32 | 0.0212 |
| **FGD1** | 1.085 | 9.03E-02 | 1.066 | 1.82E-03 | 1.248 | 2.52E-02 | -1.226 | 1.31E-06 |
| **FGD3** | 1.09 | 6.47E-01 | 1.244 | 5.77E-01 | -1.113 | 7.94E-01 | -1.026 | 6.83E-01 |
| **FGF** | -1.039 | 6.64E-01 | -1.047 | 5.69E-01 | 1.186 | 2.95E-03 | -1.133 | 1.96E-02 |
| **FilGAP** | -1.42 | 0.1649 | 1.63 | 0.1121 | -1.13 | 0.6405 | -1.05 | 0.674 |
| **FLNA** | **-1.96** | **0.0002** | **-1.51** | 0.0098 | -1.06 | 0.6852 | -1.22 | 0.4465 |
| **FN1** | 1.05 | 0.1127 | -1.11 | 0.6163 | 1.23 | 0.0367 | -1.14 | 0.0444 |
| **GIT1** | -1.16 | 0.3407 | -1.19 | 0.2232 | 1.06 | 0.7597 | -1.04 | 0.8131 |
| **GNG12** | 1.09 | 0.0156 | -1.01 | 0.0658 | 1.23 | 0.0272 | -1.11 | 0.1204 |
| **GRB2** | -1.37 | 0.0988 | -1.49 | 0.1201 | -1.12 | 0.5317 | 1.1 | 0.8963 |
| **GRLF1** | -1.09 | 0.286 | -1.15 | 0.2055 | 1.22 | 0.2017 | -1.12 | 0.4185 |
| **GSN** | **-1.62** | **0.0004** | **-1.58** | **0.1044** | 1.21 | 0.7172 | -1.25 | 0.4734 |
| **Gα12** | **1.748** | **4.30E-03** | **1.954** | **1.54E-04** | 1.239 | 6.80E-03 | 1.098 | 9.65E-02 |
| **Gα13** | **1.66** | **3.42E-05** | **1.583** | **7.92E-03** | -1.011 | 9.89E-01 | 1.06 | 8.59E-01 |
| **IQGAP** | **1.743** | **3.73E-03** | **2.096** | **5.28E-05** | 1.068 | 4.76E-01 | -1.199 | 4.12E-02 |
| **LBP** | 1.01 | 0.5477 | -1.07 | 0.8102 | 1.2 | 0.1941 | -1.11 | 0.4257 |
| **LIMK** | 1.411 | 5.13E-05 | -1.362 | 2.77E-02 | 1.158 | 7.93E-01 | 1.116 | 1.90E-06 |
| **MLCK** | -1.295 | 8.33E-02 | -1.167 | 3.21E-01 | 1.122 | 8.82E-01 | -1.236 | 1.26E-01 |
| **Myosin (MYH9)** | -1.451 | 2.52E-03 | -1.298 | 4.21E-02 | -1.076 | 7.08E-01 | -1.039 | 4.41E-01 |
| **NAP125** | 1.082 | 1.21E-02 | -1.038 | 2.90E-01 | 1.212 | 6.12E-03 | -1.142 | 2.87E-01 |
| **NHE1** | -1.08 | 0.5476 | -1.04 | 0.6144 | -1.12 | 0.5857 | 1.09 | 0.6583 |
| **NWASP** | 1.299 | 8.39E-02 | 1.361 | 4.23E-02 | -1.164 | 9.27E-02 | 1.11 | 4.92E-01 |
| **PAK** | **-2.23** | **7.87E-06** | **-3.407** | **2.49E-09** | **1.504** | **1.20E-03** | 1.016 | 6.96E-01 |
| **PDGF** | 1.434 | 1.13E-02 | **1.655** | **8.30E-05** | 1.186 | 2.93E-01 | -1.218 | 1.82E-01 |
| **PFN** | **-1.911** | **2.51E-04** | **-1.956** | **9.74E-03** | 1.179 | 4.49E-01 | 1.311 | 1.66E-01 |
| **PI3K** | **-2.088** | **2.49E-04** | **-1.932** | 1.30E-04 | -1.346 | 3.59E-01 | **1.683** | **6.09E-02** |
| **PI4P5K** | -1.206 | 1.84E-01 | -1.241 | 7.43E-02 | 1.187 | 4.52E-01 | -1.169 | 1.17E-01 |
| **PXN** | -1.14 | 0.1591 | -1.03 | 0.8886 | 1.01 | 0.7854 | -1.11 | 0.3159 |
| **RAC** | 1.249 | 1.67E-01 | 1.194 | 3.10E-01 | 1.163 | 5.00E-01 | -1.147 | 8.02E-02 |
| **RAS (RRAS2)** | **1.709** | **1.76E-02** | **1.624** | **2.58E-02** | 1.064 | 8.42E-01 | 1.107 | 4.11E-01 |
| **RHOA** | 1.46 | 0.0055 | **1.57** | **0.0002** | -1.19 | 0.9558 | 1.11 | 0.8785 |
| **ROCK** | 1.321 | 2.99E-02 | 1.314 | 1.11E-02 | 1.219 | 1.78E-01 | -1.213 | 5.32E-02 |
| **SHC** | 1.04 | 0.3897 | -1.25 | 0.0544 | 1.12 | 0.9706 | 1.1 | 0.876 |
| **SOS** | -1.498 | 6.89E-02 | -1.259 | 5.24E-01 | -1.14 | 4.76E-01 | -1.044 | 8.72E-01 |
| **SSH (SSH1)** | **-1.995** | **1.25E-05** | **-2.546** | **7.71E-04** | 1.421 | 1.85E-01 | -1.114 | 6.33E-01 |
| **Talin (TLN1)** | 1.307 | 1.48E-02 | **1.652** | **9.81E-08** | -1.146 | 3.91E-01 | -1.103 | 1.34E-01 |
| **TIAM (TIAM1)** | -1.472 | 3.66E-02 | **-1.551** | **9.84E-03** | -1.111 | 4.81E-01 | 1.171 | 3.02E-01 |
| **TMSB4** | 1.027 | 3.95E-01 | -1.067 | 7.14E-01 | 1.152 | 7.35E-01 | -1.051 | 8.66E-01 |
| **TRIO** | 1.11 | 0.3377 | 1.09 | 0.1473 | 1 | 0.4935 | 1.02 | 0.2802 |
| **VAV (VAV3)** | 1.378 | 1.33E-03 | **1.864** | **1.75E-09** | **-1.775** | **3.11E-01** | -1.27 | 1.08E-01 |
| **VCL** | -1.69 | 0.0003 | 2.25 | 0 | -1.41 | 0.0711 | -1.37 | 0.0025 |
| **WAVE1 (WASF1)** | 1.04 | 0.2366 | -1.06 | 0.7649 | 1.19 | 0.2726 | -1.11 | 0.273 |
| **WAVE2 (WASF2)** | **-1.67** | **0** | **-1.64** | **0.0001** | -1.04 | 0.8069 | 1.03 | 0.707 |
| Numbers in bold refer to each comparison with an \|fold-change\| > 1.5 and *P*<0.05 and a positive and negative fold-changes are reflected as red or green, respectively, in Figure 1. African Americans with hypertension are comparing with African American normotensives (AAHT – AANT); African Americans with hypertension compared with whites with hypertension (AAHT – WHT); African American normotensives compared with white normotensives (AANT – WNT) and whites with hypertension compared with white normotensives (WHT – WNT). | | | | | | | | |

**Supplementary Figure 1**

**AAHT vs. AANT**

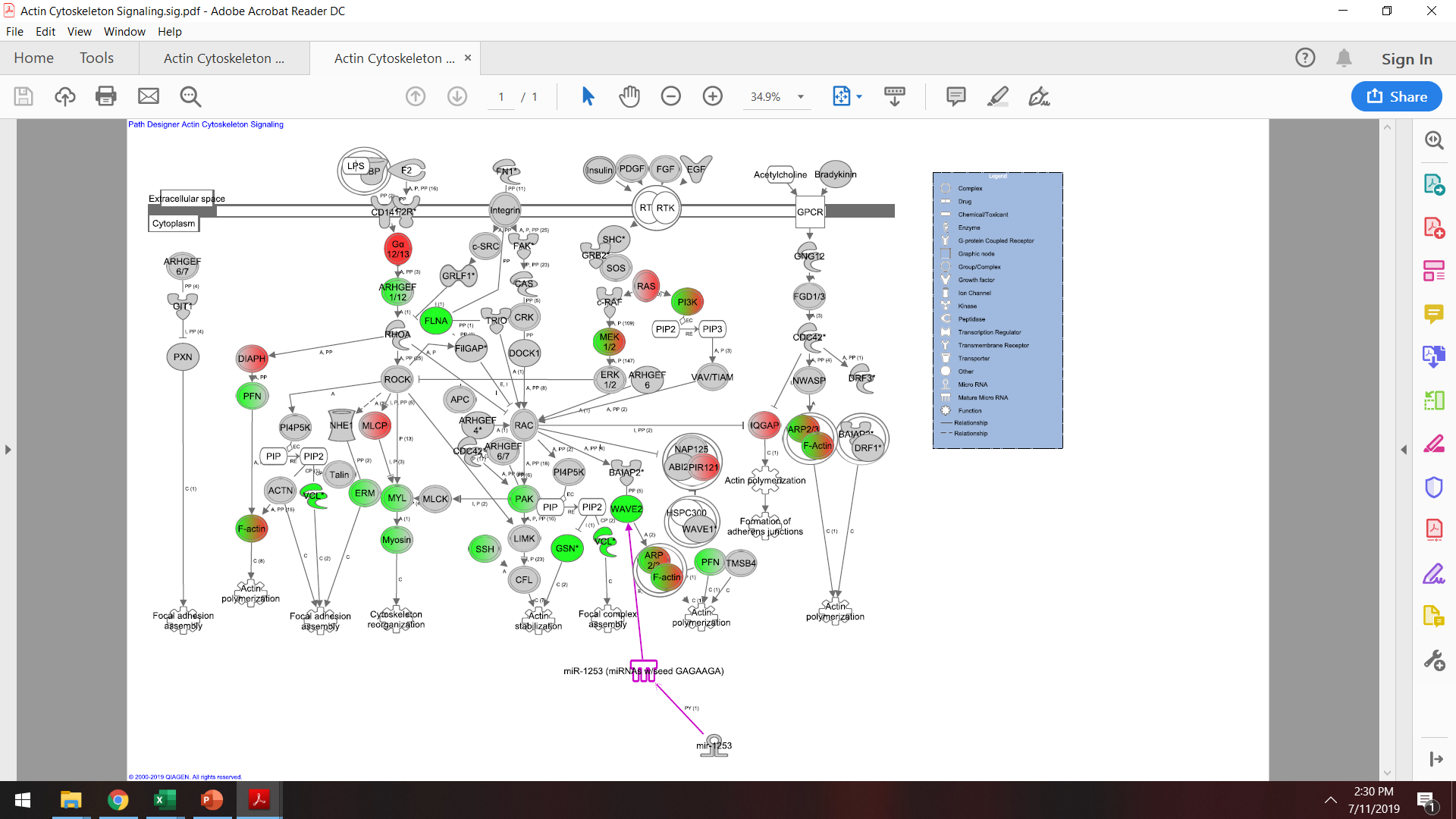

**WHT vs. WNT**

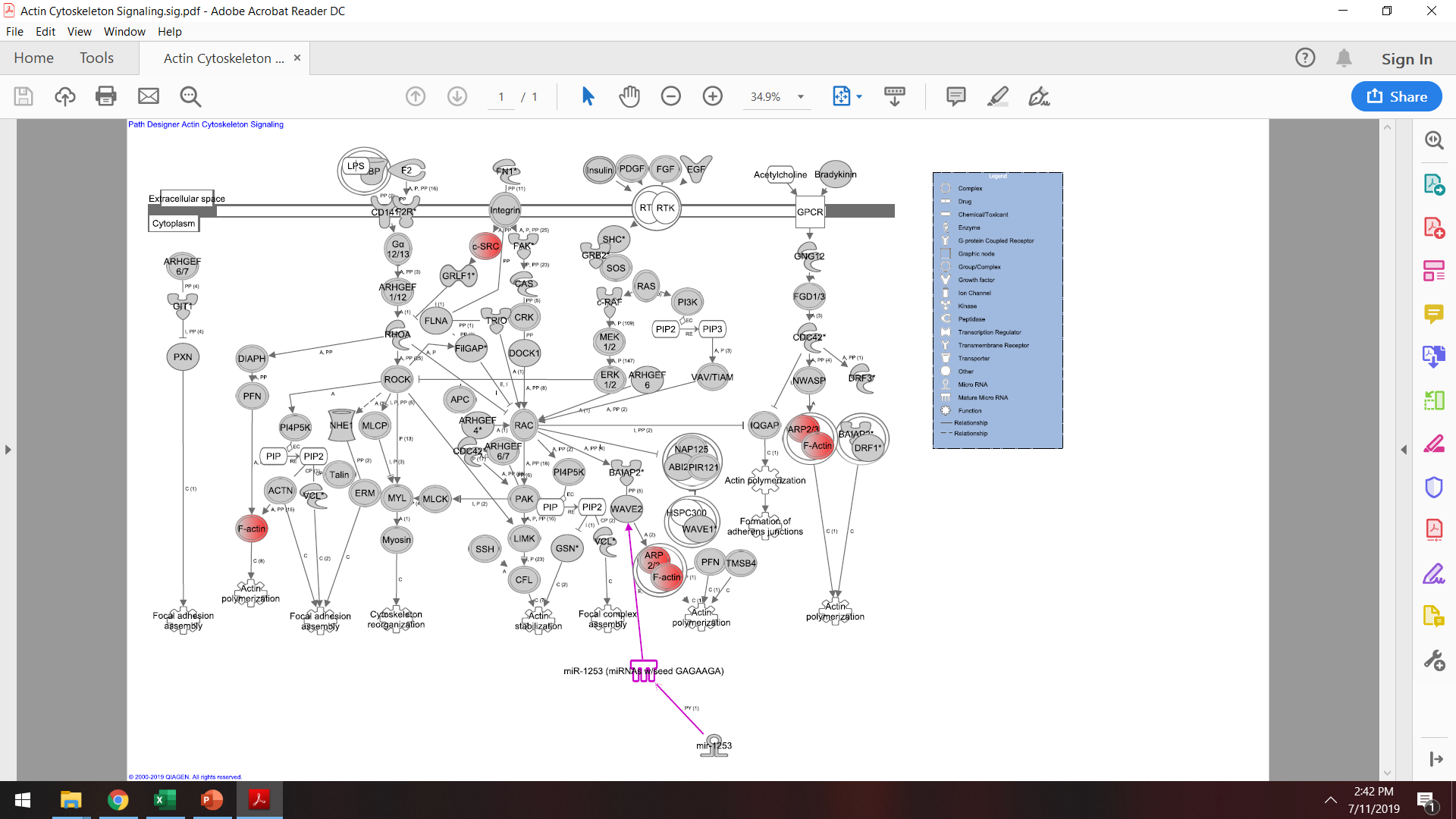

**A**

**B**

**Supplementary Figure 1: Gene expression analysis of the actin cytoskeleton in hypertensive women*.*** Microarray gene expression fold-changes in PBMCs were imported into Ingenuity Pathway Analysis (IPA) and overlaid onto the actin cytoskeleton pathway. Red indicates significantly up-regulated expression and green indicates significant down-regulation in WHT compared with WNT **(A)** and in AAHT compared with AANT **(B)**. Grey indicates a non-significant difference and white indicates no data available. All fold changes and P-values are listed for each gene and each comparison in Supplementary Table 1. AANT: Africa American normotensive; AAHT: African American hypertensive; WNT: White normotensive; WHT: White hypertensive.
